# Architecture and modular assembly of *Sulfolobus* S-layers revealed by electron cryo-tomography

**DOI:** 10.1101/677591

**Authors:** Lavinia Gambelli, Benjamin Meyer, Mathew McLaren, Kelly Sanders, Tessa E.F. Quax, Vicki Gold, Sonja-Verena Albers, Bertram Daum

## Abstract

Surface protein layers (S-layers) often form the only structural component of the archaeal cell wall and are therefore important for cell survival. S-layers have a plethora of cellular functions including maintenance of cell shape, osmotic and mechanical stability, the formation of a semi-permeable protective barrier around the cell, cell-cell interaction, as well as surface adhesion. Despite the central importance of the S-layer for archaeal life, their three-dimensional architecture is still poorly understood. Here we present the first detailed 3D electron cryo-microscopy maps of archaeal S-layers from three different *Sulfolobus* strains. We were able to pinpoint the positions and determine the structure of the two subunits SlaA and SlaB. We also present a model describing the assembly of the mature S-layer.

## Introduction

Many bacteria and archaea are surrounded by an outermost layer – the S-layer – which is composed of glycosylated surface proteins. These proteins arrange into flexible, porous, yet highly stable lattices that form cage-like coats around the plasma membrane. In bacteria, S-layers are anchored to the peptidoglycan or the outer membrane. In archaea, S-layers can be either incorporated into the periplasmatic polysaccharide layers such as pseudomurein and methanochondroitin, or simply integrated into the cytoplasmic membrane. In most cases, S-layers form ordered 2-dimensional arrays and serve a variety of functions, which are thought to be specific to genera or groups of organisms sharing the same environment (Albers & Meyer, 2011; Engelhardt, 2007; Rodrigues-Oliveira, Belmok, Vasconcellos, Schuster, & Kyaw, 2017).

For archaea, S-layers are of particular importance as they often comprise the only cell wall component. They therefore define cellular shape and provide osmotic, thermal and mechanical stability (Engelhardt, 2007). In addition, *in vitro* experiments have shown that S-layers change the physical and biochemical properties of lipid layers, rendering them less flexible, less fluid, more stable and heat-resistant, and possibly more resistant to hydrostatic pressure. Furthermore, it has been suggested that S-layers provide protection against immunological defence systems and viruses, act as pathogenic virulence factors, serve as phage receptors, promote surface adhesion, establish a quasi-periplasmic space, provide anchoring scaffolds for membrane proteins, sequester ions and facilitate biomineralization (Engelhardt, 2007). S-layers are intrinsically capable of self-assembly *in vitro*, resulting in tube-like, spherical two-dimensional crystals (Dietmar Pum & Sleytr, 2014). S-layers are therefore highly interesting for various applications in nanotechnology (Sleytr et al., 2014).

*Sulfolobus* is a genus of hyperthermophilic acidophilic archaea, which grow in low-pH terrestrial hot springs at 75–80°C all around the globe. They are well-established model organisms, as they can be relatively easily cultured in the laboratory and the genomes of a number of strains have been sequenced (Chen et al., 2005; Kawarabayasi et al., 2001; She et al., 2009). The S-layer of all *Sulfolobus* species consists of two proteins, SlaA and SlaB. Biochemical analysis of SlaA and SlaB from different *Sulfolobus* species revealed molecular masses ranging from 120kDa - 180kDa or 40kDa - 45kDa, respectively (Grogan, 2010; Veith et al., 2009). Comparative sequence analysis and molecular modelling of SlaB revealed that it exists in two species-dependent variants. In *S. ambivalens, S. acidocaldarius, S. tokodaii* and *S. sedula*, SlaB is comprised of an N-terminal Sec-dependent signal sequence, followed by three consecutive beta-sandwich domains, an alpha-helical coiled coil domain and one C-terminal transmembrane helix. In contrast, the sequences of *S. solfataricus* and *S. islandicus* are shorter by one beta-sandwich domain (Veith et al., 2009). Interestingly, it was recently shown by negative stain electron microscopy (EM) that SlaB knockout strains of *S. islandicus* still assemble partial S-layers (Zhang et al., 2018a).

In contrast, SlaA was predicted to be a soluble protein rich in beta-strands and to form the outer region of the *Sulfolobus* S-layer (Veith et al., 2009). Deletion mutants of SlaA led to deformed cells without a distinctive cell envelope (Zhang et al, 2018b).

So far, the structure of the *Sulfolobus* S-layer has only been inferred by early 2D crystallography of negatively stained and isolated S-layers. These data suggested that Sulfolobus *S-layers* adopt a structurally conserved lattice with P3 symmetry, which encompasses 4,5 nm triangular and 8 nm hexagonal pores at a 21 nm distance (Baumeister & Lembcke, 1992; Taylor, Deatherage, & Amos, 1982). However, as detailed 3D maps were so-far unavailable, it has been unclear how SlaA und SlaB assemble into the final S-layer structure.

Using electron cryo-tomography (cryoET) and sub-tomogram averaging (STA), we obtained the first 3D cryoET maps of S-layers from three *Sulfolobus* species at unprecedented resolution. Through difference maps of fully-assembled and SlaB-depleted S-layers, we were able to unambiguously pinpoint the positions of the component subunits SlaA and SlaB. Based on these experiments, we present a 3D model describing their assembly. In addition, our data reveal that strain-specific variants of SlaB lead to marked differences in the outward-facing S-layer surface. We also find that SlaB is not required for SlaA S-layer assembly *in vitro*, which has important implications about the function of both S-layer components.

## Results

### In situ structure of the *Sulfolobus* S-layer

To obtain a first detailed understanding of the molecular architecture of the archaeal S-layer, we chose the *Sulfolobus* species *S. islandicus (Sisl), S. solfataricus (Ssol)* and *S. acidocaldarius (Saci)*. In *Sisl* and *Ssol* the S-layer subunit proteins SlaA and SlaB show share 87.4% and 87.7% sequence identity (Fig. 1S1) and are thus virtually identical. In contrast, SlaA and SlaB from *Saci* show only ∼24% or ∼25 % identity with *Sisl / Ssol* (Fig. 1S2) and are thus far less conserved between those species. Moreover, SlaB from *Saci and Sisl/Ssol* have previously been proposed to represent two different structural “families” with respect to SlaB, which exists as a long form in *Saci* and a short form in *Ssol* and *Sisl* (Veith et al., 2009). We hypothesised that the two different variants of SlaA and SlaB may cause distinct S-layer geometries in *Saci* when compared to *Sisl* or *Ssol*.

**Figure 1.**
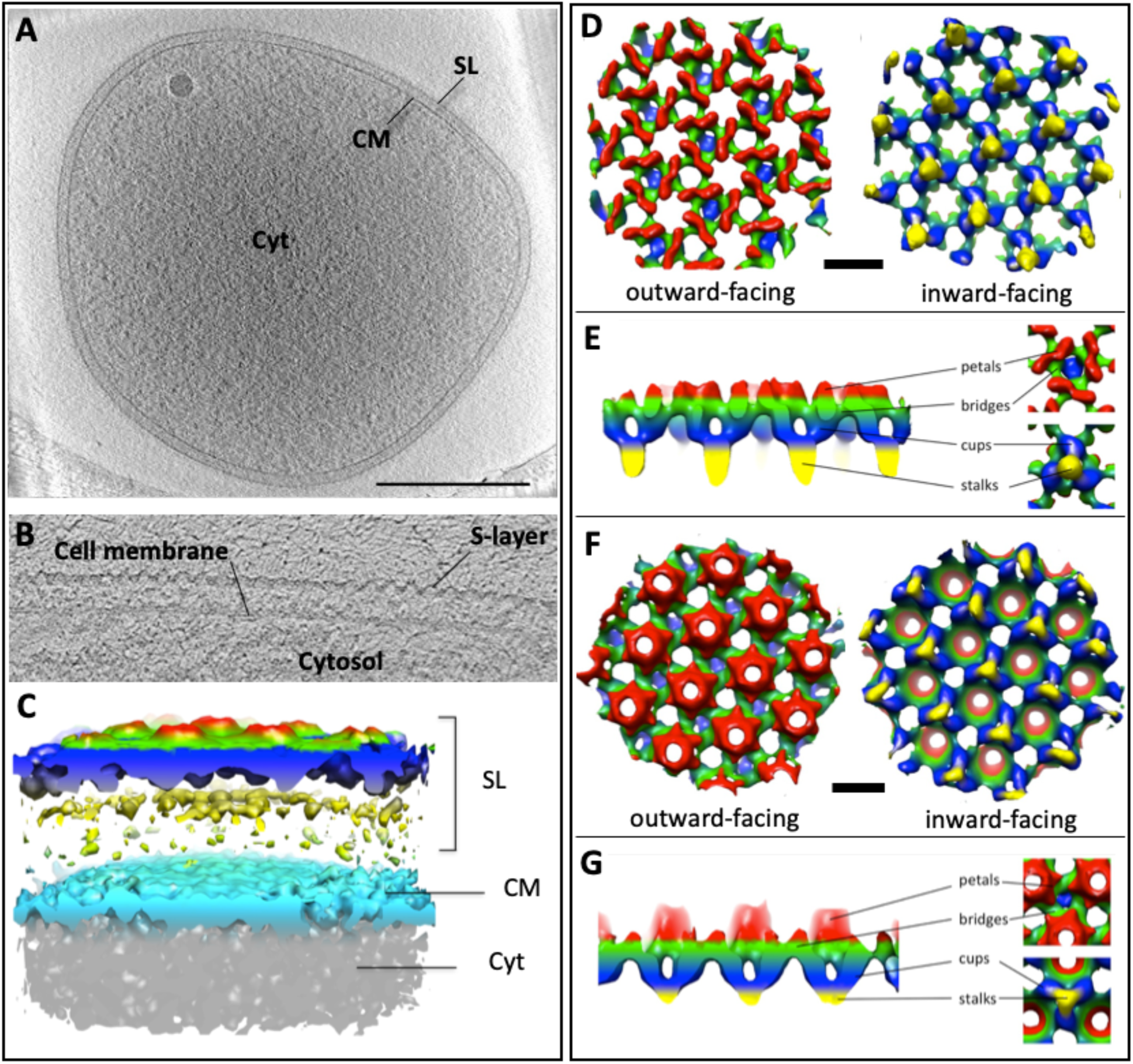
Cryo electron tomography of *Sulfolobus* cells. **A**, tomographic slice through a *Sisl* cell. **B**, cross section through the cell membrane and S-layer. **C**, segmented surface representation of the *Sisl* S-layer. Scale bar (A), 500 nm. **D-G**, Sub-tomogram averages of *Saci* (D,E) and Sisl (F,G) S-layers. Maps are coloured by proximity to the membrane plane. Yellow, proximal; red, distal. Scale bars (D, F), 20 nm.

To prepare cells for cryoET, cellular suspensions were plunge-frozen on holy carbon grids and investigated in the electron microscope. The majority of cells were surrounded by intact S-layers and membranes and showed various degrees of cytoplasmic density, which may either be a result of different metabolic states of slight cytosolic leakage during the sample preparation procedure. Tomographic tilt series were collected of cells with low cytosolic density, which were more transparent to the electron beam and thus resulted in tomograms with better signal-to-noise ratio (Fig. 1A).

In tomographic reconstructions, cells appeared disk-shaped with a diameter of up to 2 μm and a thickness of 250-300 nm (Fig. 1A). Since *Sulfolobus* cells are usually roughly spherical, this suggests that the cells had been compressed due to surface tension of the buffer during the plunge freezing procedure. In tomographic cross-sections, two layers confining the cells were distinguished (Fig. 1A). Whereas the inner layer, the membrane, was smooth, the outer S-layer had a corrugated appearance (Fig. 1B).

Tomographic sections parallel to the plane of the S-layers clearly showed that they indeed form regular two-dimensional arrays, as confirmed by power spectra of the respective tomographic slices. These power spectra showed clear spots up to the third order and indicated a lattice with P3 symmetry (Fig. 1S3). The fuzzy spots indicated that the S-layers were not perfectly crystalline. This was expected, as roughly round shapes cannot be contained in a hexagonal lattice unless defects are included (D Pum, Messner, & Sleytr, 1991). Moreover, gaps in the lattice are needed to accommodate surface filaments such as archaella (Daum et al., 2017) or to allow the cells to grow and divide. In tomographic slices perpendicular to the membrane plane, S-layers formed regular, corrugated canopy-like arrays at a centre-to-centre distance of ∼30 nm from the membrane (Fig. 1 B).

To obtain structural information of these S-layers in situ, 1809 subvolumes (*Sisl*) and 2068 subvolumes (*Saci)* were cut from tomograms, in which the cell surface was clearly resolved. Subsequently, subvolumes were aligned and averaged in PEET. This resulted in 3D maps at ∼30 Å and 28 Å resolution for *Sisl* and *Saci*, respectively (Fig. 1S5). Both S-layer maps revealed a perforated two-dimensional protein lattice with P3 symmetry (Fig. 1; 1S4; 1S5). For *Sisl*, unit cell dimensions were ∼21.9 × 20.9 nm including an angle of 120° and for *Saci* the unit cell measured ∼23.9 × 23.6 nm with an angle of 120°. The unit cell of the *Saci* S-layer is therefore roughly 10 % larger than that of *Sisl.* The total height of the maps was ∼25 nm for *Sisl* and ∼28 nm for *Saci*, respectively (Fig. 1 D-G).

Our 3D maps revealed structurally conserved and unique features for both S-layers (Fig. 1 D-G). In particular, the organisation of the inward-facing domains of the S-layer of both species are generally the same. In either case, they consist of trimeric stalks (Fig. 1 blue and yellow). These stalks support an outer canopy-like assembly (Fig. 1 D-G, green and red). At their membrane-proximal parts, each of the trimeric stalks merge into a single protrusion, giving them a tulip-like shape (Fig. 1 E, G blue and yellow). This protrusion most likely forms the membrane anchor of the S-layer. Distal to the membrane, the stalks are connected to the S-layer canopy via their trimeric domains.

The outer canopy forms the outward-facing side of the S-layer and resides on top of the array of the trimeric stalks (Fig. 1 D-G). In *Sisl* and *Saci* the base of the canopy appears to be cross-linked by a protein network, which has roughly the same in-plane geometry in both species (Fig. 1 D-G, green). In cross-section perpendicular to plane of the S-layers, this network forms a continuous band throughout the entire structure and includes repeating triangular and hexagonal pores. Two types of triangular pores can be distinguished, of which one is situated on top of a trimeric stalk (Fig. 1, E, G blue) while the other is not. The triangular pores with and without stalk alternate throughout the lattice (Fig. 1 D-G).

The outward-facing part of the canopy is distinct in both species. In *Sisl*, it is composed of an array of hexameric cone-shaped protrusions (Fig. 1 G, red). Each of the cones encloses a hexagonal pore within the membrane. At their base, each cone is surrounded by six neighbouring cones. The spaces between three cones contain the triangular pores (Fig 1 G).

In *Saci*, the most distal part of the outer layer is composed of elongated, petal-like structures (Fig. 1 D, red). As in *Saci* these structures do not protrude as far from the general plane of the canopy, the outer S-layer face appears rather smooth compared to that of *Sisl*. As seen in Fig. 1 D, each of these structures show a 2-fold symmetry around their long axis, which suggests that they are dimers.

### Structural dissection of *Sulfolobus* S-layers

To obtain higher resolution maps for *Sulfolobus* S-layers and to investigate if the cell context is important to maintain correct S-layer assembly, we isolated S-layers – this time from *Saci* and *Ssol*, the latter of which was by sequence alignments predicted to have the same shortened domain structure as *Sisl.* Tomograms were recorded at optimised imaging conditions and sub-tomogram averaging resulted in more detailed maps (Fig. 2) at 21 Å (*Saci*) and 16 Å (*Ssol*) resolution (Fig. 1S5). The lattice parameters of the *Saci* S-layer remained unchanged, showing that their integrity does not crucially depend on the cellular context (Fig. 2 A). Moreover, the map of the *Ssol* S-layer was virtually identical to the one of *Sisl* in terms of lattice dimensions and outer canopy topology (Fig. 2 B), indicating that 87 % sequence identity conserves the general structural features.

**Figure 2.**
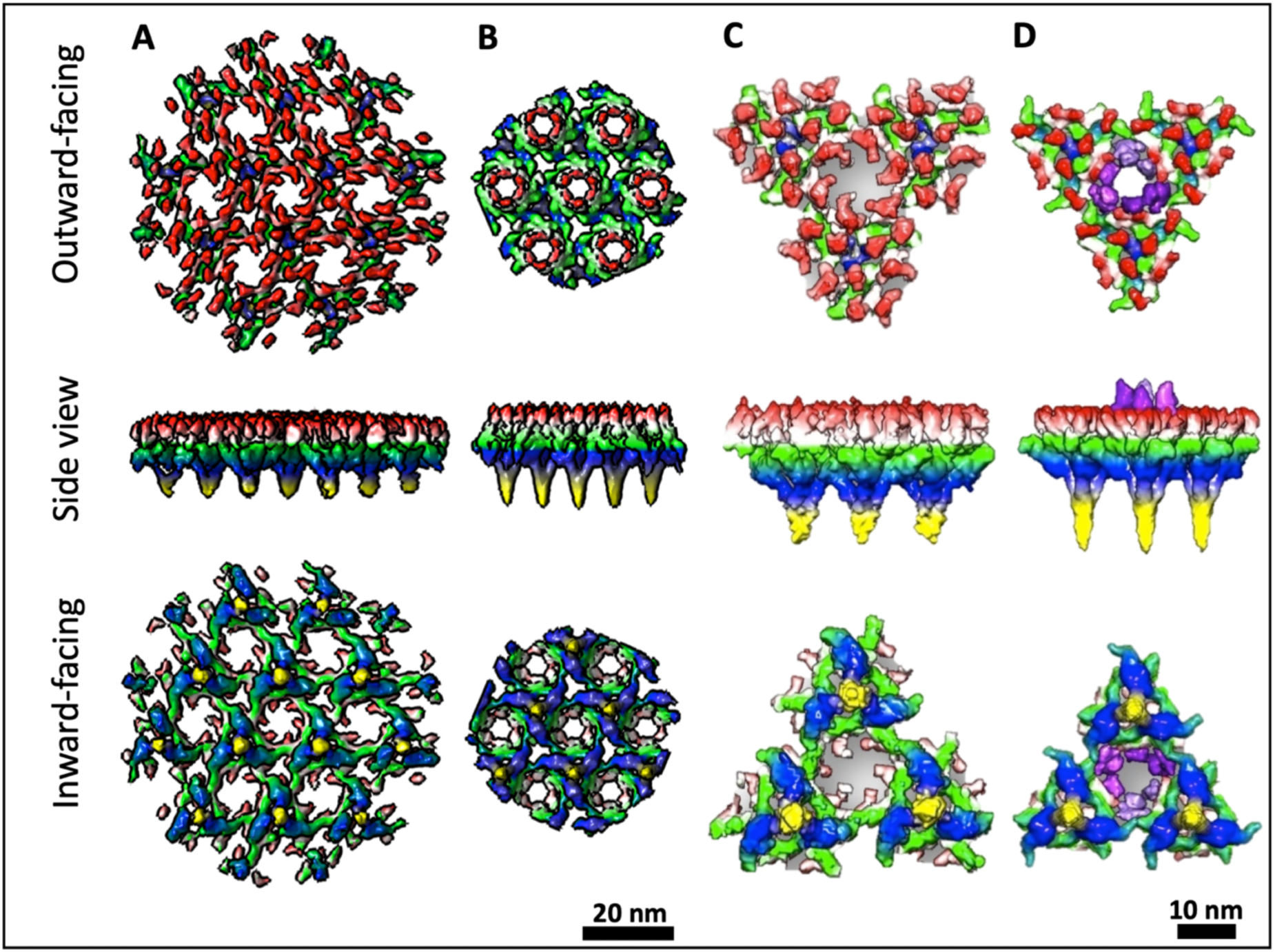
Structure of isolated S-layers from *S. acidocaldarius* and *S. solfataricus*. **A, B**, S-layer maps of *Saci* (A) and *Ssol* (B) at 21 Å or 16 Å resolution, respectively. **C, D**, S-layer unit of *Saci* (C) and *Ssol* (D), including three stalks and a hexameric pore. Maps are coloured by proximity to the membrane plane. Yellow, proximal; red, distal; purple, conical protrusions. Scale bars, 20 nm.

To further investigate the structural differences between *Saci* and *Ssol*, we segmented each map into units composed of three stalks and the adjacent canopy (Fig 2 C, D). As with *Sisl*, the S-layer of *Ssol* clearly showed a smaller unit size of 10 % when compared to the one of *Saci*. Furthermore, structural differences surrounding the hexagonal pores were resolved more clearly. Whereas in *Saci* each hexagonal pore is flanked by six inward-curled protein domains, it is surrounded by six outward projecting densities in *Ssol*, giving rise to the cone-like assemblies.

### Pinpointing SlaA and SlaB

In order to locate SlaA and SlaB within the *Sulfolobus* S-layer, we first removed the subunit SlaB from isolated S-layers using the detergent N-laurylsarcosine (Fig. 3 A). After 3-4 repeated washing steps, this subunit ceased to be detectable by SDS PAGE. CryoET and sub-tomogram averaging of the SlaB-depleted S-layer resulted in a map that lacked the trimeric stalks (Fig 3 C), leaving behind the porous canopy. This clearly indicates that SlaB constitutes the trimeric stalks that anchor the S-layer in the membrane. Consequently, the canopy must be formed by SlaA, which is in line with previous predictions (Veith et al., 2009).

**Figure 3.**
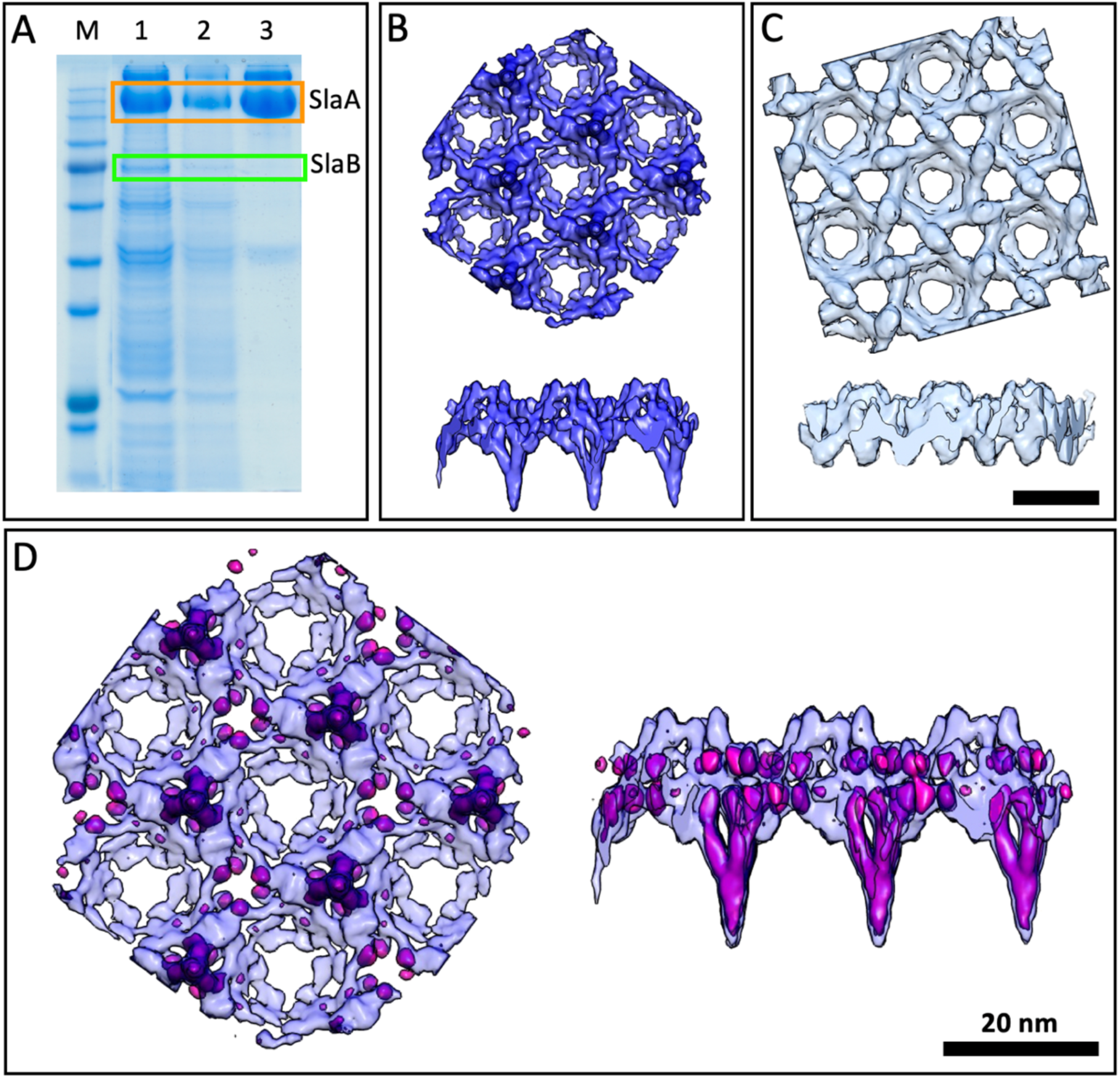
Structural dissection of the *S. solfataricus* S-layer. **A**, SDS PAGE. M, marker; 1, washed once; 2, washed twice; 3, watched three times in detergent. **B**, sub-tomogram average of fully-assembled S-layer. **C**, sub-tomogram average of SlaB-depleted S-layer. **D**, difference map (pink) overlayed with the complete S-layer visualises location of SlaB.

To visualise the location and architecture of SlaB unambiguously, we calculated a difference map by subtracting the SlaB-depleted from the fully assembled map (Fig. 3 D). This revealed that SlaB adopts a tripod-like shape, which is consistent with earlier sequence-based predictions that suggested that SlaB foms a trimer (Veith et al., 2009). In our structure, the three branches of each SlaB trimer are buried inside the SlaA canopy, whilst the “monomeric” stalk projects away from it, towards the membrane plane. In multitude, these pillars act to raise the SlaA canopy above the membrane. Interestingly, individual SlaB units are not in contact, but are linked to each other via the SlaA canopy network. Notably, the canopy structure of the SlaB-depleted S-layer was less well resolved than in the fully-assembled control sample (Fig. 3 B). This suggests that SlaB may act in reinforcing the stability of the SlaA network.

### Role of SlaB in S-layer assembly

As the S-layer structure is maintained after removal of SlaB, we asked if SlaB proteins merely function as pillars for the SlaA lattice or if they also aide correct assembly of the outer canopy. To investigate this, we isolated S-layers from both *Saci* and *Ssol*, disassembled them by transfer into pH 10 buffer and subsequently performed recrystallisation by dialysis against water at pH 7 and incubation for 120 h. Inspection of the reaction containing both subunits in the electron microscope revealed patches of reassembled S-layers (Fig. 4 A, D). Sub-tomogram averaging showed that these S-layers had retained their original lattice dimensions (Fig. 4 B, E) and that the SlaA canopy as well as SlaB stalks could be identified (Fig. 4 C, F).

**Figure 4.**
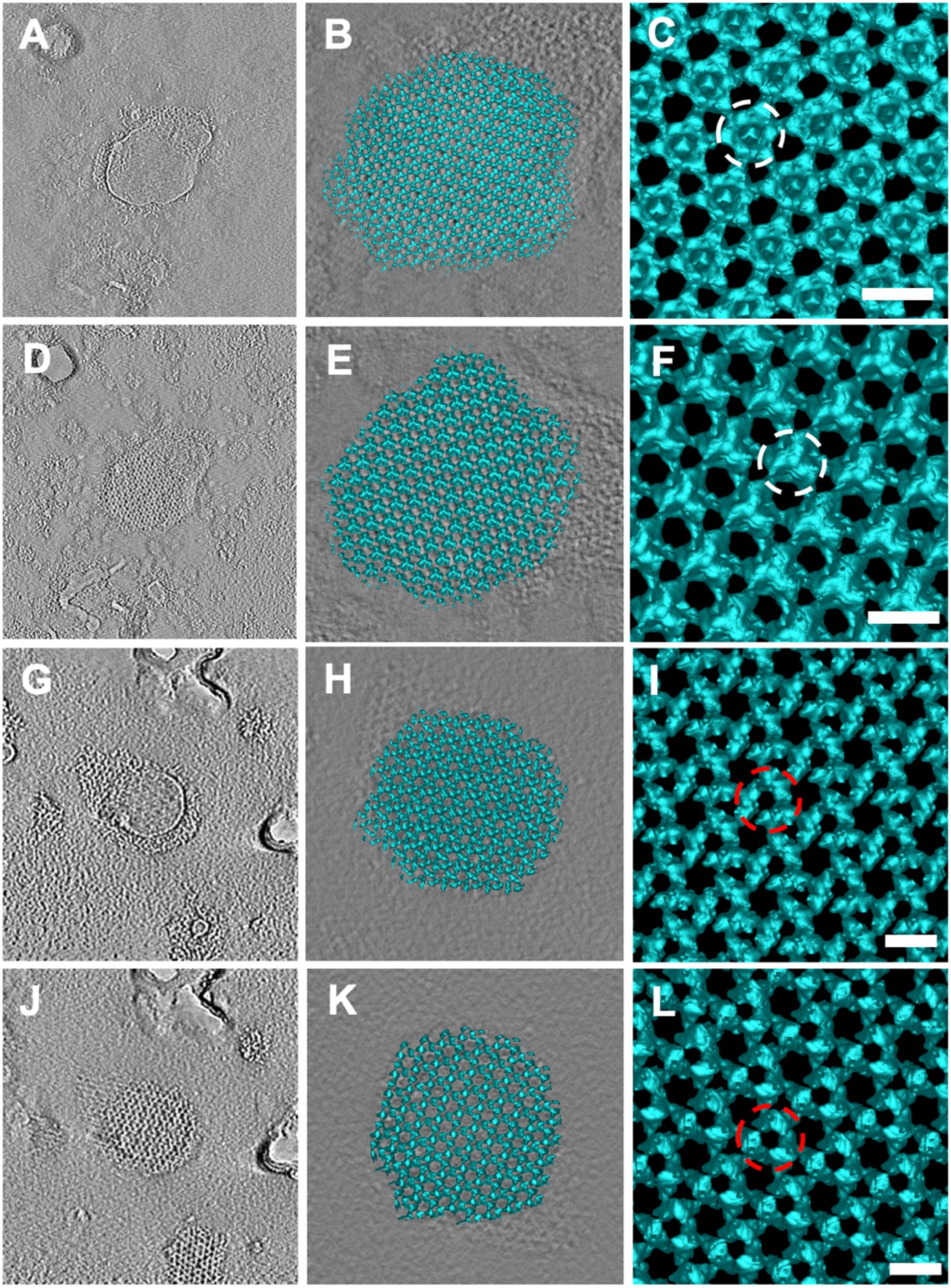
S-layers recrystallise with and without SlaB. A-F, reassembly of *Sulfolobus* S-layers from SlaA and SlaB. G-L, reassembly of S-layers from SlaA only. (A,D,G,J, tomographic slices; B,E,H,K, 3D surface representation; C,F,I,L, sub-tomogram averages). White circle highlights presence of SlaB (C&F) and red circles absence of SlaB (I&L). Scale bar: 20 nm

Interestingly, electron microscopy of the SlaB-depleted samples also revealed newly-formed S-layers (Fig. 4 G, H, J, K). Again, sub-tomogram averaging confirmed that the reassembled S-layers had adopted their original lattice parameters, however this time without the SlaB stalks (Fig. 4 I and L). These observations indicate that SlaB is not involved in assembly process of the SlaA canopy and suggests that it mainly functions as a membrane anchor and distance ruler that determines the width of the periplasmic space.

### Assembly model of the *Sulfolobus* S-layer

It has previously been shown biochemically and confirmed by us that SlaA forms a stable ∼250 kDa dimer in solution (Veith et al., 2009). In conjunction with early electron microscopy, it has also been predicted that this dimer forms the primary *Sulfolobus* S-layer building block (Taylor et al., 1982; Veith et al., 2009). This repeating dimer-array is apparent in our sub-tomogram averages of negatively-stained SlaA-only S-layers (Fig. 4 I). To build a cryoEM model of the *Saci* and *Ssol* S-layers, we first segmented the SlaB-depleted sub-tomogram averaging maps by the water-shedding method in Chimera (Pettersen et al., 2004). For guidance, we used the constraints of repeating dimeric SlaA densities that roughly fitted the molecular weight of 250 kDa, as well as our negative stain map in which the individual dimers were clearly visible (Fig. 4 I). We then superimposed this segmentation with the fully-assembled S-layer, which provided us with the positions of SlaB trimers.

The resulting model shows the location of SlaA dimers and SlaB trimers within the assembled S-layers of *Saci* (Fig. 5 A-C) and *Ssol* (Fig. 5 D-F). In both cases, individual SlaA dimers adopt the shape of a boomerang. The elbow of each boomerang interacts with one of the three branches of a SlaB trimer, whereas both SlaA arms project outward. Thus, each SlaB trimer is bound to three SlaA dimers (SlaB_3_/3SlaA_2_), which intersect to form a triangular pore above each SlaB trimer. Three of these flower-shaped SlaB_3_/3SlaA_2_ surround one hexagonal pore, formed by six SlaA dimers. Herein, each of the three SlaB trimers contributes 2 SlaA dimers to the hexameric ring. At the same time, each SlaA dimer spans two neighbouring hexameric pores. The second triangular pore type that does not coincide with a SlaB trimer is formed by three intersecting SlaA dimers at the interface of three hexagonal pores. In total, each SlaA dimer interacts with four SlaA dimers and 3 SlaB trimers. Therefore, SlaA dimers interlink into a tightly woven S-layer canopy that is stabilised by multiple interactions.

**Figure 5.**
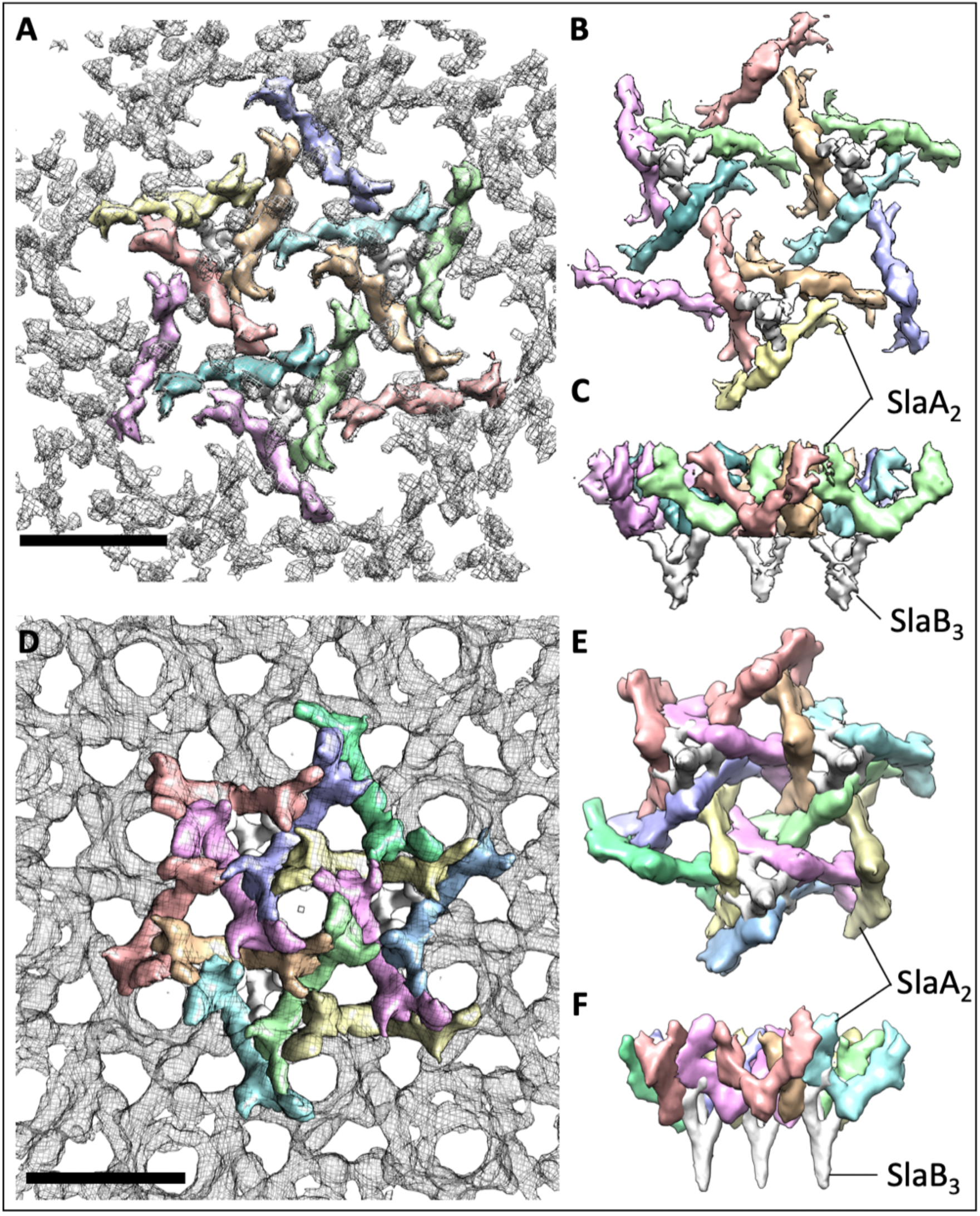
Assembly models for the *S. acidocaldarius* (A-C) and *S. solfataricus* (D-F) S-layers. **A, D** outward facing surfaces (grey mesh) segmented into SlaA dimers (each dimer in a different colour) and SlaB trimers (white). **B, E** inward-facing surface. **C, F** side view, perpendicular to the membrane plane. Scale bar: 20 nm

### SlaA determines S-layer geometry and topology

To understand how *Saci* and *Ssol* assemble S-layers with similar geometry but different topology and unit cell size, we compared the structures of the SlaA proteins from each species. To this end, we extracted individual dimer densities from our segmented maps shown in Fig. 5.

This comparison revealed that while the overall shape of the dimers is the same, there are marked differences with respect to their horizontal and vertical dimensions, as well as the angles between both arms of each boomerang (Fig 6). *Saci* SlaA measures 23 nm along its long axis, which corresponds with the size of the unit cell of the S-layer (Fig 6A). This is to be expected, as this molecule spans to neighbouring hexagonal S-layer pores (Fig. 6C). The corresponding portion of *Ssol* SlaA measures ∼20 nm (Fig. 6B), which is again in accordance with the 10% smaller assembled array (Fig 6D). The length of the SlaA dimer therefore determines the size of the unit cell. The differences in length appear to be established by the angle between both arms of the molecule, which is 102° for the longer *Saci* SlaA and 96° for the shorter *Ssol* homolog. Interestingly, both SlaA variants also differ in an apical domain, which is curled inwards in *Saci* and projects upwards in *Ssol*. In assembly, these domains are responsible for the differences in the topology of both S-layers, as it is these domains that delineate the shallow hexagonal pores in S*aci* and the conical protrusions in *Ssol* (Fig. 6 C, D). Taken together, the strain-specific dimensions and surface morphology of *Sulfolobus* S-layers are determined by unique structural characteristics of SlaA, in particular its apical domains.

**Figure 6.**
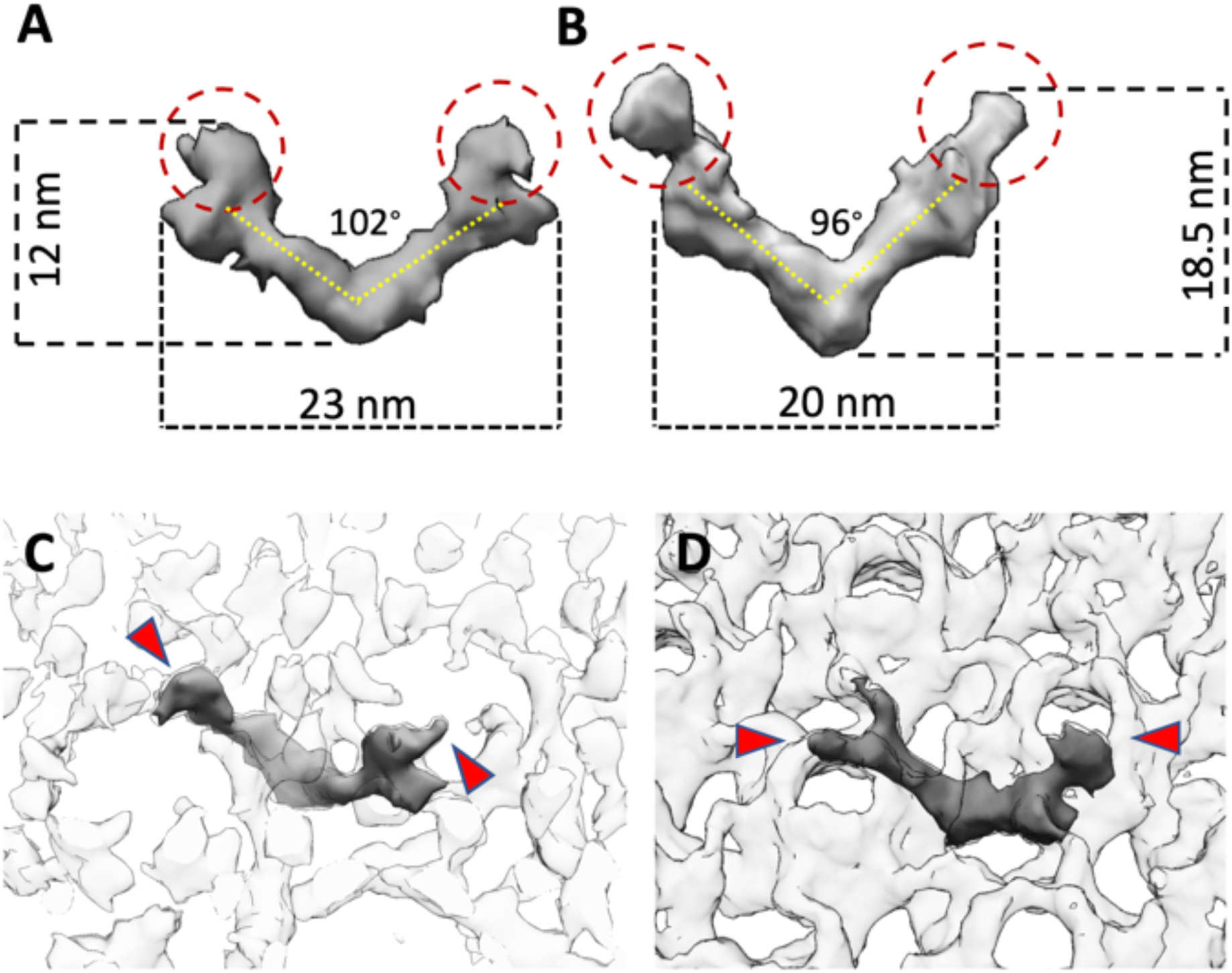
SlaA structure determines S-layer topology and unit dimensions. Comparison of the structures of SlaA from *Saci* (**A**) and *Ssol* (**B**) show differences in length, height and angular shape of the dimer. **C, D**, location of one SlaA dimer within the S-layer of *Saci* (**C)** and *Ssol* (**D**). Red circles / arrowheads indicate apical domains that determine shape and topology of the hexameric pores.

## Discussion

### Roles of SlaA and SlaB in S-layer assembly

In this study, we describe the first cryoET-based molecular maps of archaeal S-layers. These maps unambiguously show that the *Sulfolobus* S-layer consists of trimeric SlaB proteins that adopt tripod-like structures and act as pillars that anchor the outer S-layer canopy in the cell membrane. The building block of the S-layer canopy is a dimer of SlaA, which assembles into a porous array of P3 symmetry. These observations are consistent with earlier predictions based on sequence analysis, biochemistry and early negative-stain electron microscopy maps (Baumeister & Lembcke, 1992; Taylor et al., 1982; Veith et al., 2009).

While X-ray structures for neither of the component proteins are available, sequence predictions indicated that SlaA and SlaB contain Sec-dependent signal peptides and are thus translocated through the membrane by the general secretory pathway. In addition, glycoproteomic analysis showed that both proteins contain several sites for N- and O-linked glycosylation and are heavily glycosylated prior to assembly (Palmieri et al., 2013; Peyfoon et al., 2010) (Fig. 7). Moreover, SlaB has been predicted to consist of an N-terminal membrane-anchor, followed by coiled-coil domain including a serine and threonine-rich highly-glycosylated region (ST linker) and 2-3 C-terminal beta sandwich domains (Veith et al., 2009) (Fig. 7). Upon trimerisation of SlaB, three C-termini presumably form a trimeric coiled-coil, which most likely corresponds to the stalk-like base of each SlaB tripod (Fig. 7). The SlaB stalks in our maps appear shorter than expected for a coiled-coil of roughly 100 amino acids (Veith et al., 2009) in length. This is likely due to high flexibility in this region, which is thus mostly averaged out during our sub-tomogram averaging procedure. The three protrusions of each SlaB tripod are likely formed by the predicted C-terminal beta-sandwich domains (Veith et al., 2009) of the three SlaB proteins in the trimer, which therefore also form the interface with the overarching SlaA lattice (Fig. 7). However due to the limited resolution of our maps, we were unable to distinguish between the length differences in the *Saci* and *Ssol* SlaB beta-sandwich domains.

**Figure 7.**
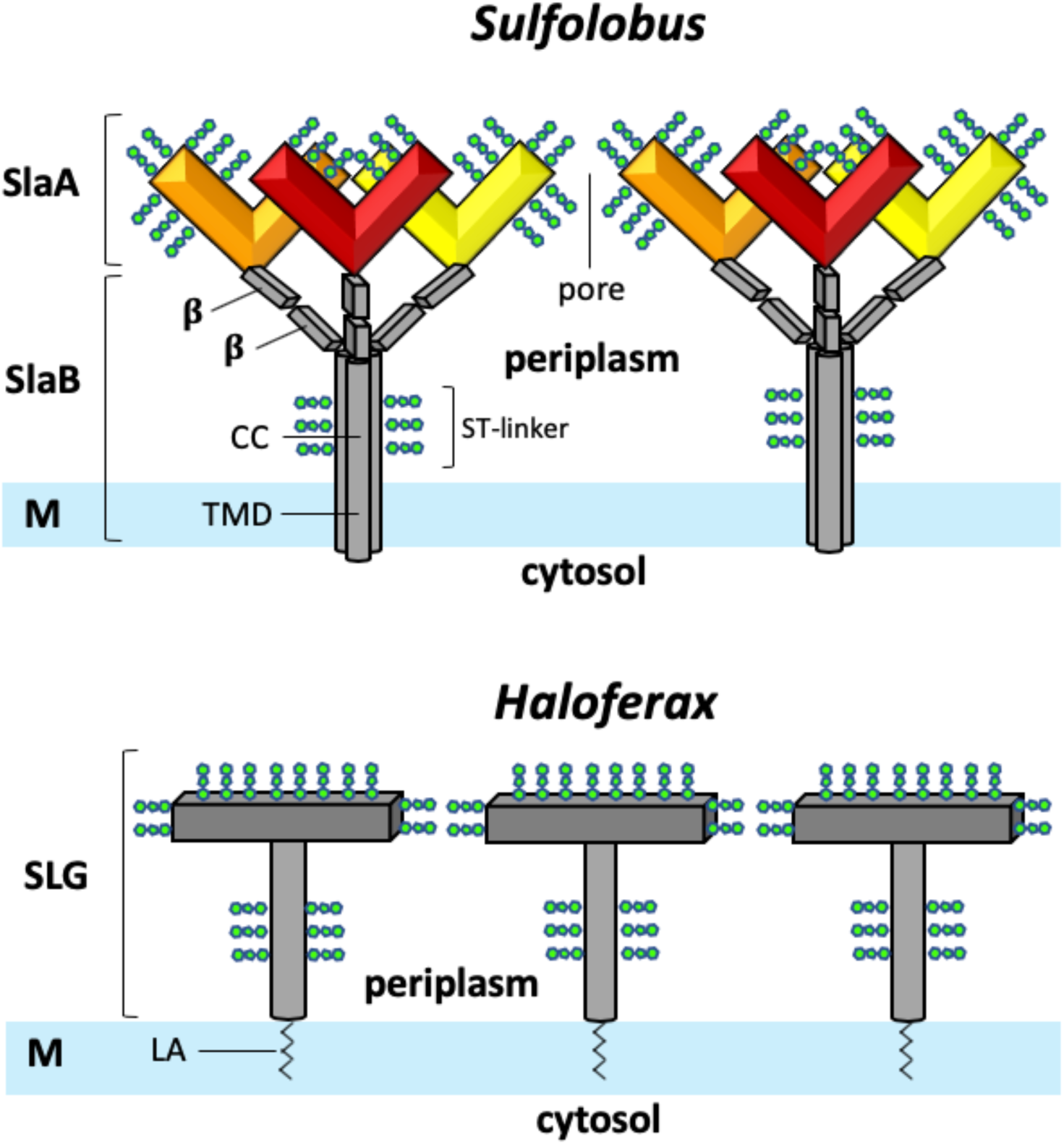
Model of the *Sulfolobus* S-layer compared to the single-subunit S-layer of haloarchaea. The Sulfolobus S-layer (upper panel) consists of the two protein subunits: SlaA dimers (red, orange, yellow) form the outer S-layer canopy. Each SlaA protein is predicted to be rich in β strands. The SlaA dimer has a boomerang-like shape, the angle of which determines the S-layer unit cell size. SlaB trimers (grey) form the membrane anchors of the S-layer. Each SlaB is predicted to consist of an N-terminal transmembrane domain (TMD), a coiled-coil domain (CC) and 2-3 C-terminal β-sandwich domains (β). SlaA and SlaB proteins are highly glycosylated (green). In contrast, the S-layers of many halophile archaea (such as that of Haloferax volcanii; bottom panel) consist of multiple copies of only one S-layer glycoprotein (SLG) subunit and are inserted into the membrane by a posttranslationally added lipid anchor (LA).

Our reassembly experiments reveal that SlaB is not required for SlaA self-assembly *in vitro*. This is consistent with recent whole-cell negative stain EM data, which showed that *Sisl* SlaB knockout strains still assemble partial S-layer like coats (Zhang et al., 2018a). We therefore conclude that the SlaA protein is the driving factor of S-layer assembly. In addition, our data shows that strain-specific S-layer surface features including lattice dimensions and topology, are determined by the shape of SlaA.

In contrast, SlaB’s main function is to serve as a membrane anchor, distance ruler and to create the periplasmic space. This space has been proposed to have important roles in archaeal cell biology, including the coordination of membrane-embedded molecular machines, such as the archaellum (Banerjee et al., 2015; Daum et al., 2017) and various other archaeal surface structures (Chaudhury, Quax, & Albers, 2018). Secondarily, we observe that SlaA-only S-layers are slightly less ordered than their fully-assembled counterparts, indicating that SlaB also increases the stability of the S-layer, presumably by crosslinking 3SlaA_2_ subcomplexes with each other.

The functional significance of the different S-layer topologies is largely unknown and awaits further exploration. It may be speculated that distinct S-layer surface structures have evolved to modulate the cell-adhesive properties of different archaeal strains. In addition, S-layers may act as receptors in cell-cell recognition or phage infection (Plaut et al., 2014) and we could show previously that archaeal viruses evolved elaborate cell entry and egress strategies to overcome the S-layer barrier (Daum et al, 2013; Quemin et al, 2014; Quax & Daum 2018). It is therefore conceivable that distinct S-layer topologies provide unique recognition tags for strain-specific interaction and communication and that the evolution of new structures is a manifestation of the cell’s strategy to avoid viral infection (Tschitschko et al, 2015). SlaB, in contrast, appears structurally more conserved than SlaA. Indeed, it has been shown previously that SlaB proteins have lower sequence variability than SlaA and their molecular masses differ less across different *Sulfolobus* strains (Veith et al., 2009). This is likely due to the fact that SlaB interacts less with the extracellular medium and is thus less prone to environmentally-related evolutionary pressures.

Recently, an additional paralogous SlaB gene product (M164_1049, SSO1175 or WP_011278654.1 in *Sisl, Ssol*, or *Saci*, respectively) was suggested to form another membrane-anchored protein component of the *Sulfolobus* S-layer (Zhang et al, 2018b). Double M164_1049 and SlaB mutants showed that no S-layer is present in *Sisl*, as opposed to single SlaB mutants where a partial coat of SlaA was observed. In addition, a single M164_1049 deletion mutant showed virtually no difference compared to the wild type. It was concluded that M164_1049 may aid in anchoring SlaA to the cytoplasmic membrane (Zhang et al., 2018b). Based on our maps, we cannot confirm the presence of this protein. As it is membrane-anchored, it likely resides near the membrane plane, the region which is averaged out in our data due to the highly flexible nature of the SlaB coiled-coil domain.

In contrast to *Sulfolobus*, many archaeal strains possess only one S-layer protein. For example, in many haloarchaea, a single S-layer glycoprotein (gene product *csg1* / SLG protein in *Haloferax volcanii*) combines membrane-anchoring and canopy-forming functions into one protein. Moreover, this protein is not membrane-integral but instead inserted to the bilayer via a C-terminal lipid anchor (Fig. 7) (Pohlschroder, Pfeiffer, Schulze, & Halim, 2018). While employing only one S-layer protein might be energetically more favorable, it is likely that using two increases the adaptability of the S-layer surface (SlaA) without compromising the membrane anchor (SlaB).

### Function of S-layer pores

One of the most striking features of the *Saci* and *Ssol* S-layers are the circular apertures of ∼4.5 or ∼8 nm in diameter that repeat throughout the array. Similar porous, skeleton-like architectures have been reported for other S-layers such as that of the bacterium *Caulobacter cresentus* (Bharat et al., 2017), highlighting that S-layer porosity is a highly conserved trait throughout bacterial and archaeal species and thus provides a significant evolutionary advantage. This poses the question what function these openings may convey. It is likely that – similar to chainmail - this skeleton-like structure enhances S-layer flexibility, enabling it to coat cell bodies of various shape and allowing the cell to morph and divide. It is also tantalising to assume that the particular shape and size of the pores has evolved to accommodate particular cellular filaments, such as adhesive pili or archaella. In contrast to this notion, we find that most known archaeal surface filaments, such as adhesive (AAP) pili (Braun et al., 2016; Henche et al., 2012) and archaella (Daum et al., 2017; Poweleit et al., 2016) are over 10 nm in diameter and thus too wide to be accommodated by S-layer pores. Thus, the S-layer will have to be partially disassembled or adopt a different local geometry wherever these filaments emerge from the cell body (Daum et al., 2017). Finally, it is safe to assume that due to their porosity, S-layers provide a semi-permeable barrier, similar to the outer membrane of Gram-negative bacteria, albeit with a more liberal molecular weight cut-off. At the first glance, this cut-off appears to be determined by the pore diameter (<8nm), which would allow a large variety of solutes, macromolecules and even large proteins to pass. However, this notion may be deceptive, as both SlaA and SlaB are highly glycosylated (Palmieri et al., 2013; Peyfoon et al., 2010). While (likely due to their high flexibility) these posttranslational modifications are averaged out in our cryoEM maps, they are thought to cover much of the S-layer surface and thus possibly also project into the pores (Fig. 7). It is conceivable that these glycans would significantly lower the permeability of the S-layer pores to macromolecules, similar to the hydrogel found in nuclear pore complexes (D’Angelo & Hetzer, 2008).

## Conclusions

We present the first detailed 3D models of S-layers of three different *Sulfolobus* strains and pinpoint the location and organisation of their component subunits SlaA and SlaB. We find that the structure of the SlaA dimer determines the unit cell size and topology of the S-layer, whilst SlaB anchors the S-layer in the plasma membrane and defines a pseudo-periplasmic space. Different S-layer topologies between species are likely the result of various evolutionary adaptations, including cell-cell interactive specificity and adhesive properties. Despite being an ubiquitous hallmark in archaea, many functional aspects of S-layers still need to be clarified through further structural investigation.

## Materials and Methods

### Strains and growth conditions

The strains *Ssol* and *Saci* MW001 were grown in basal Brock medium* at pH 3 (Brock, 1972). The medium was supplemented with 0.1 % (w/v) NZ-amine and 0.2 % (w/v) dextrin just before inoculation. For growth of *Saci* MW001, 10 μg/ml uracil was added to the medium. *Sulfolobus* cultures were grown at 75°C, 150 rpm, until an OD600 of > 0.6 was reached. Cells were harvested by centrifugation at 5000 x g (Sorvall ST 8R) for 30 min and stored at -20°C for subsequent use.

* Brock media contains (amount per litre): 1.3 g (NH_4_)_2_SO_4_, 0.28 g KH_2_PO_4_, 0.25 g MgSO_4_ × 7H_2_O, 0.07 g CaCl_2_ × 2H_2_O, 0.02 g FeCl_2_ × 4H_2_O, 1.8 mg MnCl_2_ × 4H_2_O, 4.5 mg Na_2_B_4_O_7_ × 10H_2_O, 0.22 mg ZnSO_4_ × 7H_2_O, 0.05 mg CuCl_2_ × 2H_2_O, 0.03 mg NaMoO_4_ × 2H_2_O, 0.03 mg VOSO_4_ × 2H_2_O and 0.01 mg CoSO_4_ × 7H_2_O.

### S-layer isolation

Cell pellets of frozen cells from a 50 ml culture were incubated and inverted on a rotator at 40 rpm (Stuart SB3) for 45 min at 37°C, in 40 ml of buffer A (10 mM NaCl, 1 mM phenylmethylsulfonyl fluoride, 0.5 % sodium lauroylsarcosine), with the addition of 10 μg/ml DNAse I just prior to use. The samples were pelleted by centrifugation at 18.000 x g (Sorvall Legend XTR) for 30 min, and subsequently resuspended in 1.5 ml buffer A, before further incubation and inversion at 37°C, for 30 min. After centrifugation at 14.000 rpm for 30 min (Sorvall ST 8R), the pellet was purified by resuspension and incubation in 1.5 ml buffer B (10 mM NaCl, 0.5 mM MgSO_4_, 0.5% SDS) and rotated for 20 min at 37°C, 40 rpm. To retain both SlaA and SlaB, only one wash was performed. To remove SlaB, washing with buffer B was repeated a further three times. Purified S-layer proteins were washed once with distilled water and stored at 4°C. The removal of SlaB was confirmed by SDS-PAGE analysis.

### Disassembly and reassembly of S-layers

S-layers were disassembled by increasing the pH with the addition of 20 mM NaCO_3_, pH 10, 10 mM CaCl_2_ and incubation at 60°C, 600 rpm (Thermomixer F1.5, Eppendorf) for 1h. Disassembly was verified by SDS-PAGE analysis.

Reassembly was achieved by buffer exchange using 0.5 ml concentrators according to the manufacturer’s instructions (Pierce), with dH_2_O to reduce the pH to 7, followed by the addition of 10mM CaCl_2_ and incubation at 60°C for 120 hours.

### Negative stain tomography

A 3 μl sample of S-layers with and without SlaB was placed on glow-discharged carbon-coated copper grids (Agar Scientific) and incubated at room temperature for 1 min. The excess sample was removed by blotting using filter paper (Whatman, GH healthcare). The specimens were stained using ammonium molybdate for 1 min. Grids were blotted dry with filter paper and let air dry completely. Single axis tilt series (−60 to +60) were recorded on a FEI T12 electron microscope operated at 120 kV, using DigitalMicrograph (Gatan Inc.).

Tilt series were reconstructed using the IMOD package (Kremer et al, 1996) and tomograms were generated using the weighted back-projection algorithm.

### Electron cryo-tomography

3 µl of S-layer suspension were applied to glow-discharged 300 mesh copper Quantifoil grids (R2/2, Quantifoil, Jena, Germany), blotted for 3 – 5 seconds and rapidly injected into liquid ethane using a homemade plunge-freezer. Tomograms were recorded using a Polara G2 Tecnai (Thermo-Fisher, Eindhoven, The Netherlands) or a Jeol 3200 FSC TEM (JEOL, Tokyo, Japan), both based at the Max Planck Institute of Biophysics in Frankfurt, Germany. The microscopes were operated at 300 kV. Data were collected using a 4 × 4 k CCD camera or a 4×4 k K2 Summit direct electron detector (Gatan Inc., Pleasanton, USA) running in counting mode. Inelastically scattered electrons were removed using a Gatan Tridiem energy filter (Gatan Inc., Pleasanton, USA) for the Polara and an in-column energy filter for the JEOL microscope. Tilt series were collected in zero-loss mode using Digital Micrograph (Gatan, Pleasonton, USA) from -62° to +62° and in steps of 2°. For tomograms of whole cells, the Polara was used. The magnification was set to 41.000 × (resulting in a final pixel size of 5.4 Å on the final image) and defocus values of 6-8 μm were applied. For isolated S-layers, the magnification used was 72.000 (final image pixel of 2.8 Å) on the Polara and 10.000 x (final pixel size 3.35) on the JEOL. Defocus values of 2-4 μm were used. Whole-cell tomograms were recorded using a CCD camera with a total dose of 100 e^-^ Å^-2^. Tomograms of isolated S-layers were recorded using the K2 and in dose-fractionation mode at a dose rate of 8-10 e^-^ px^-1^ s^-1^ and a maximum total dose of 60 e^-^ Å^-2^. Dose-fractionated tilt images were aligned using an in-house script based on IMOD (Kremer et al, 1996) programmes, CTF corrected and reconstructed into tomograms using the IMOD software package (Kremer et al, 1996).

### Sub-tomogram averaging

To average the S-layer, a random grid of points was applied over the S-layer within a tomograms of *Sulfolobus* cells or isolated S-layers, which were previously binned twofold. Whole-cell tomograms were filtered by NAD (Frangakis and Hegerl, 2001) to enhance the contrast. Using PEET (Nicastro et al, 2006; Heumann et al, 2011), subvolumes were extracted, aligned and averaged. For the *in situ* S-layer average, 1809 subvolumes (*Sisl*) and 2068 subvolumes (*Saci)* of 60×60×60 pixels in dimension were used. For isolated S-layers, 325 subvolumes (*Saci)* and 5561 subvolumes (*Ssol)* of 100 × 100 × 80 were used. P3 symmety was applied to the final averages. S-layers were visualised and segmented using UCSF Chimera (Pettersen et al, 2004). The resolution of all averages was estimated based the reflections in their respective power spectra calculated by IMOD (Kremer et al, 1996). For the *in situ* structures this suggested a resolution of 28 Å for *Saci* and 30 Å for *Sisl*. For the maps of the isolated S-layers, 16 Å were measured for *Ssol* and 21 Å for *Saci* (Fig. 1S5).

### Difference maps and assembly models

The difference maps were calculated and assembly models built in UCSF Chimera (Pettersen et al, 2004). To calculate the difference map between the fully assembled and SlaB-depleted S-layers, the *vop subtract* command was used. This resulted in clear densities for SlaB trimers. To build the assembly model of the SlaA canopy regions, semi-automated water-shedding segmentation was initially used on the sub-tomogram average of the SlaB-depleted S-layers. This approach segmented the map in equal-sized parts, which were then grouped to generate a P3 lattice of dimers. Individual SlaB and SlaA subunits were extracted as separate densities from the map and fitted into the fully assembled *Ssol and Saci* S-layers to yield the complete SlaA/SlaB assembly model.

### Sequence alignments

Protein sequences were sourced online using Uniprot (https://www.uniprot.org/) and multiple sequence alignments performed with the Praline Server (http://www.ibi.vu.nl/programs/pralinewww/)

## Acknowledgements

We thank Werner Kühlbrandt and Deryck Mills (MPI of Biophysics in Germany) for allowing us to use the Polara electron microscope. This project was funded by the Max Planck Society (BD), the University of Exeter Research Fellow’s Startup grant (BD) and the ERC starting grant “MICROROBOTS” (BD, LG, MM, KS).

---

**Figure 1 S1.**
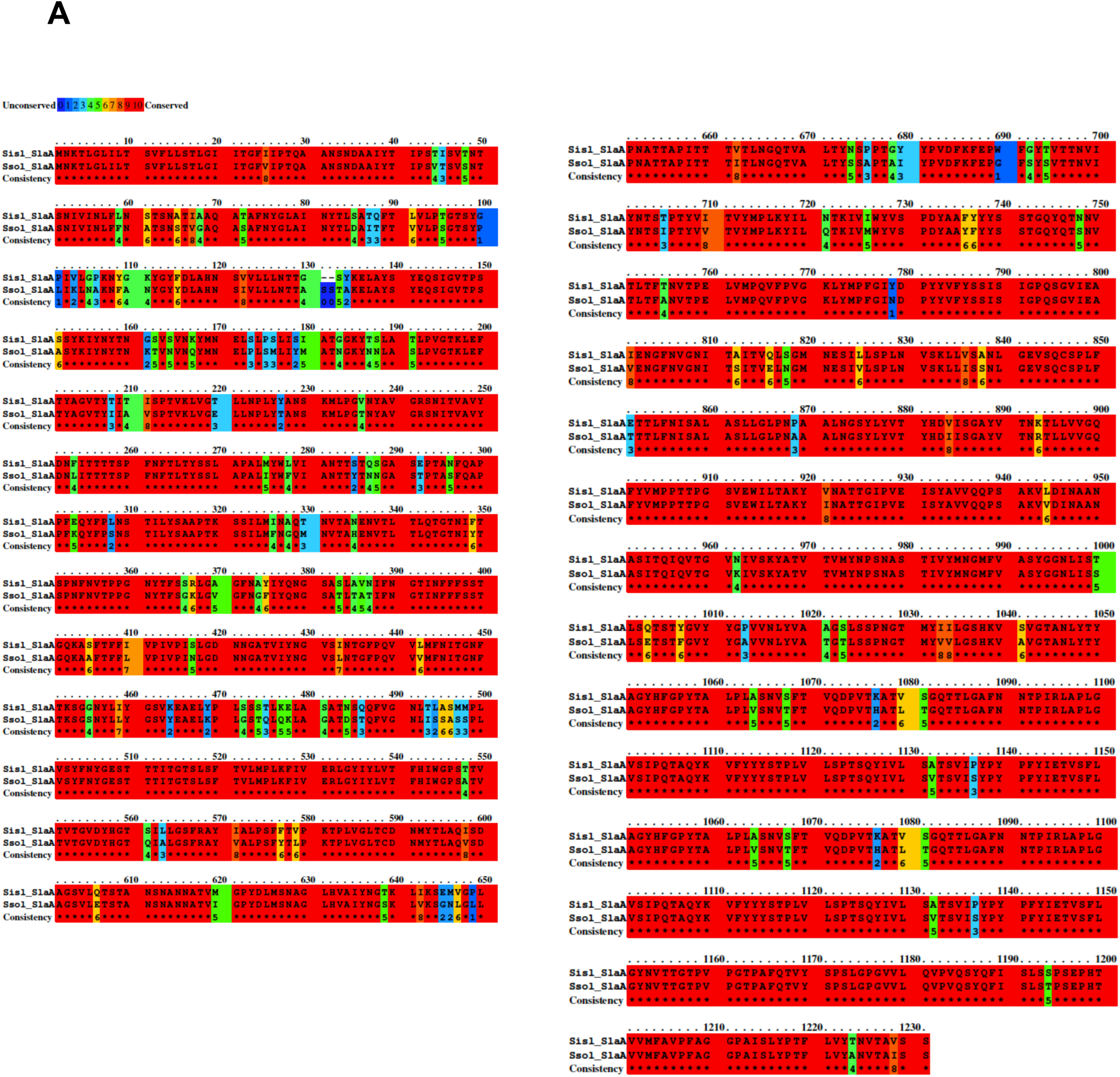
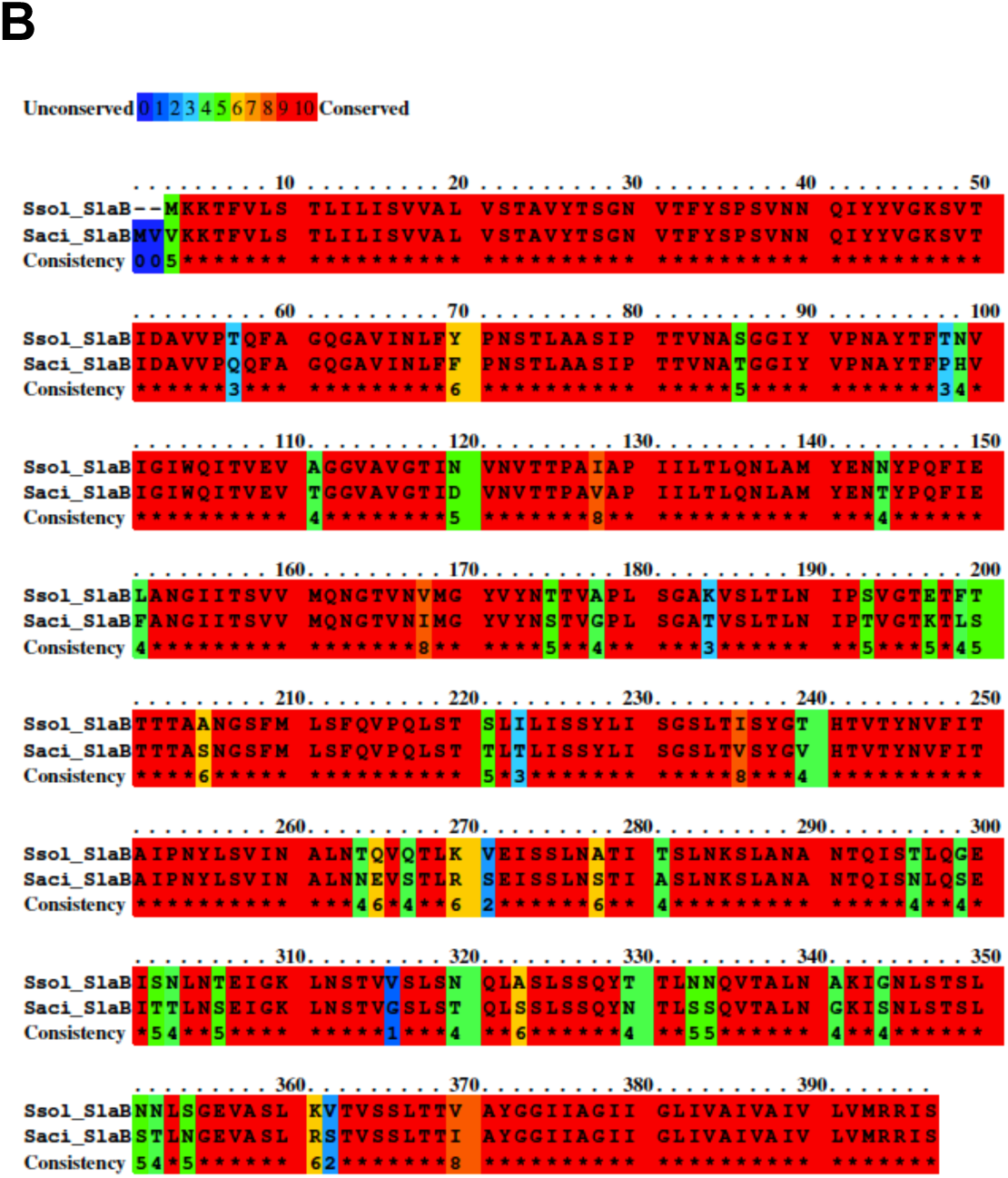
Sequence alignment comparing SlaA (A) and SlaB (B) from *Sisl* and *Ssol*. The sequence identity for SlaA and SlaB is 87.4% and 87.7%, respectively.

**Figure 1 S2.**
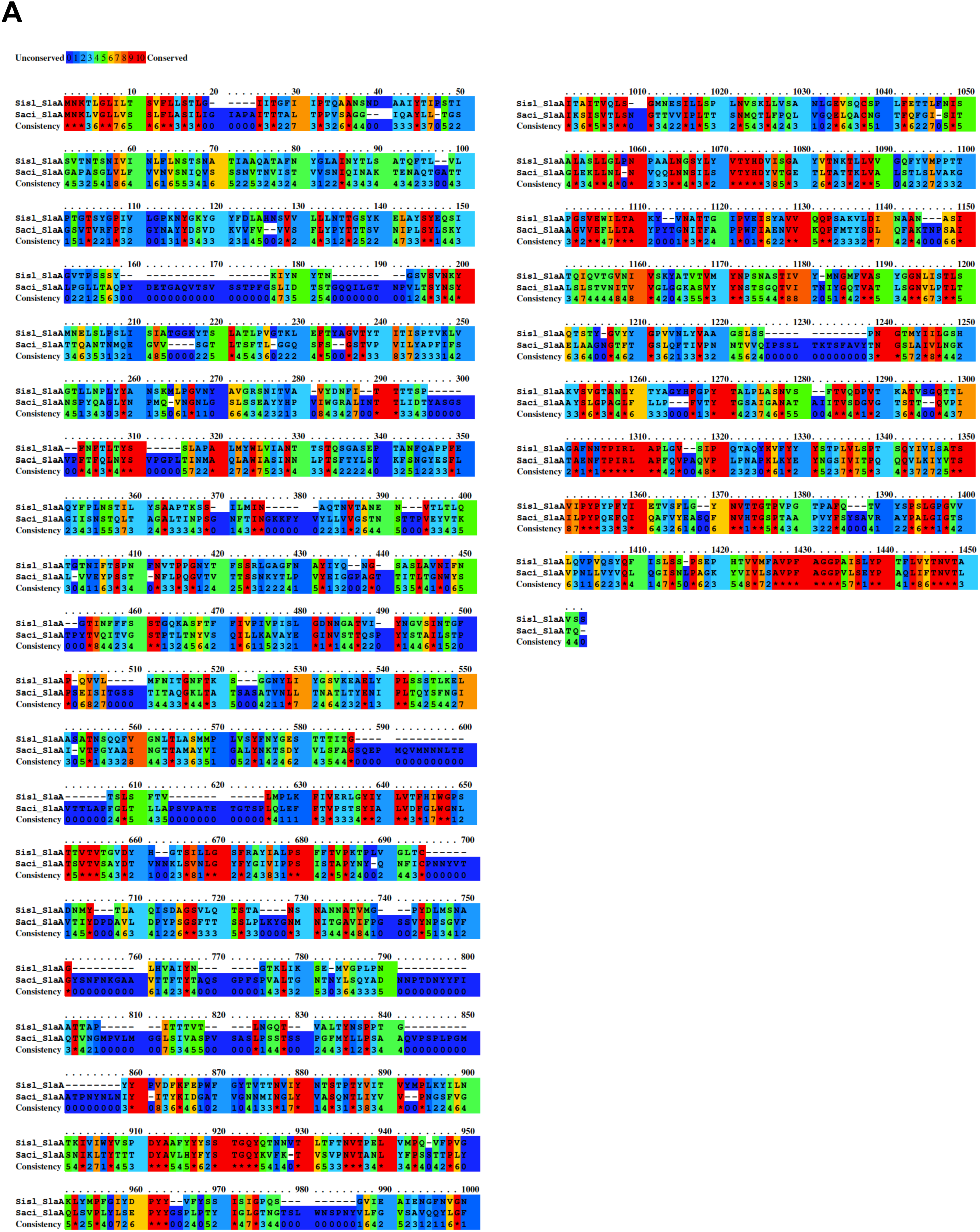
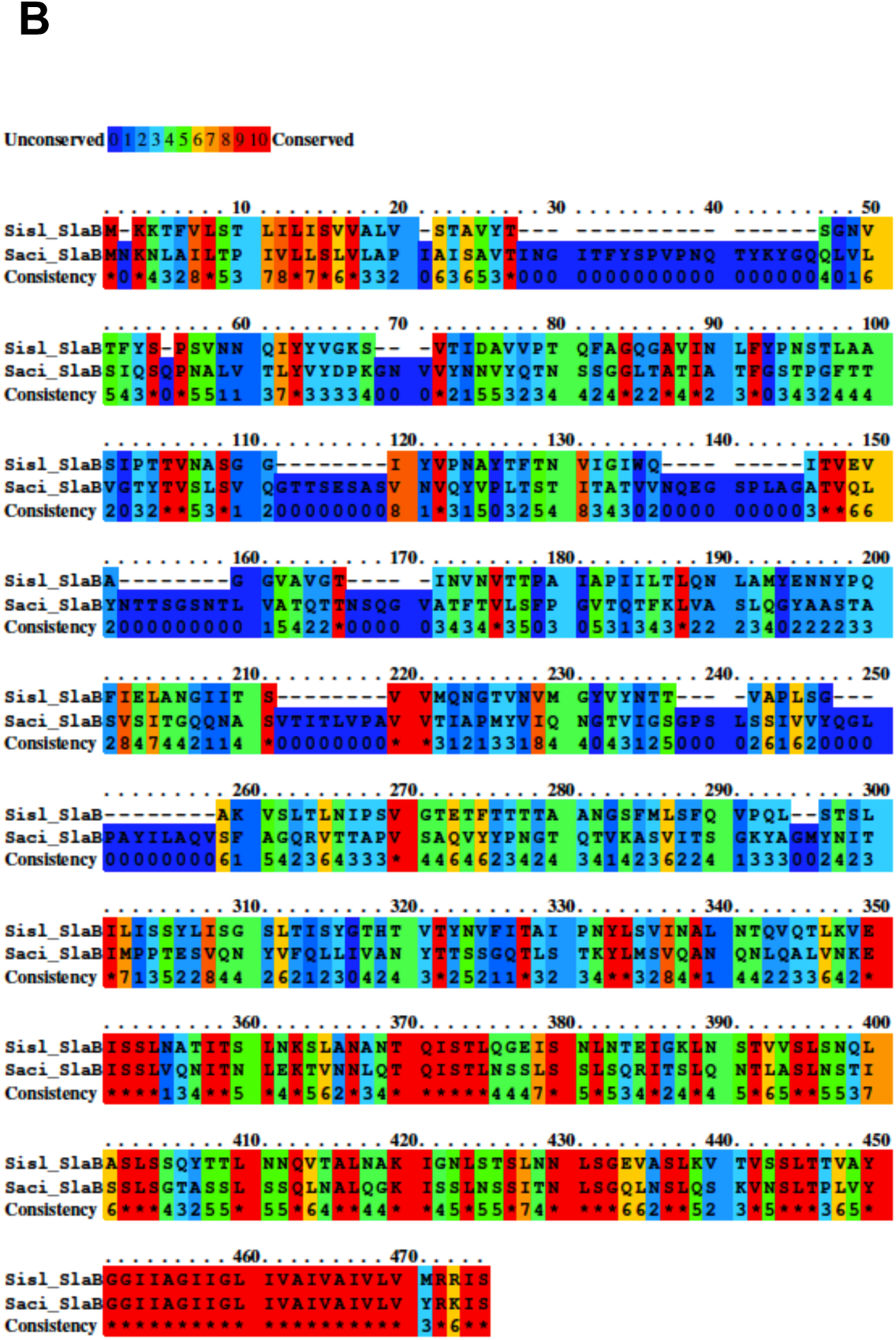
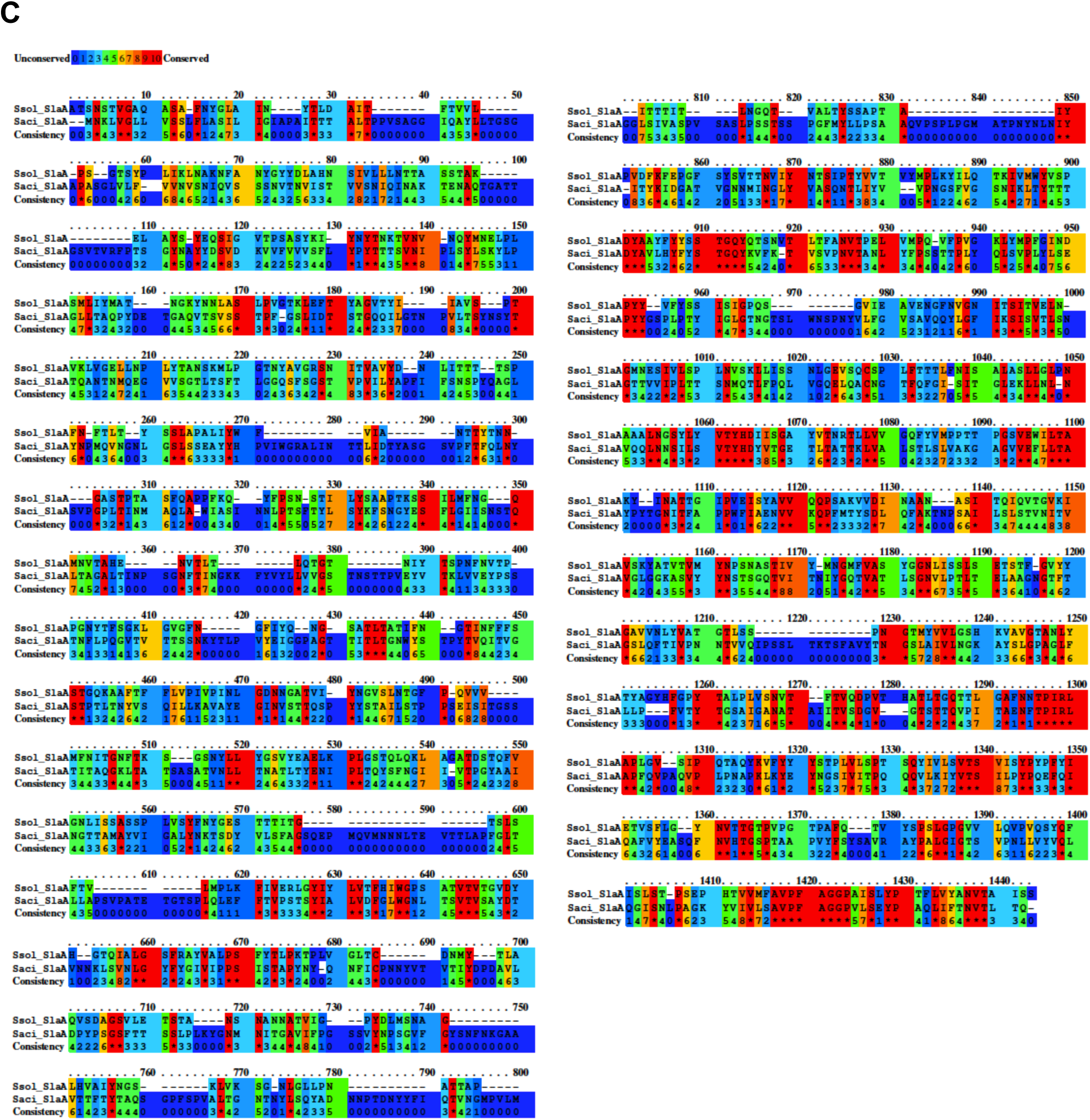
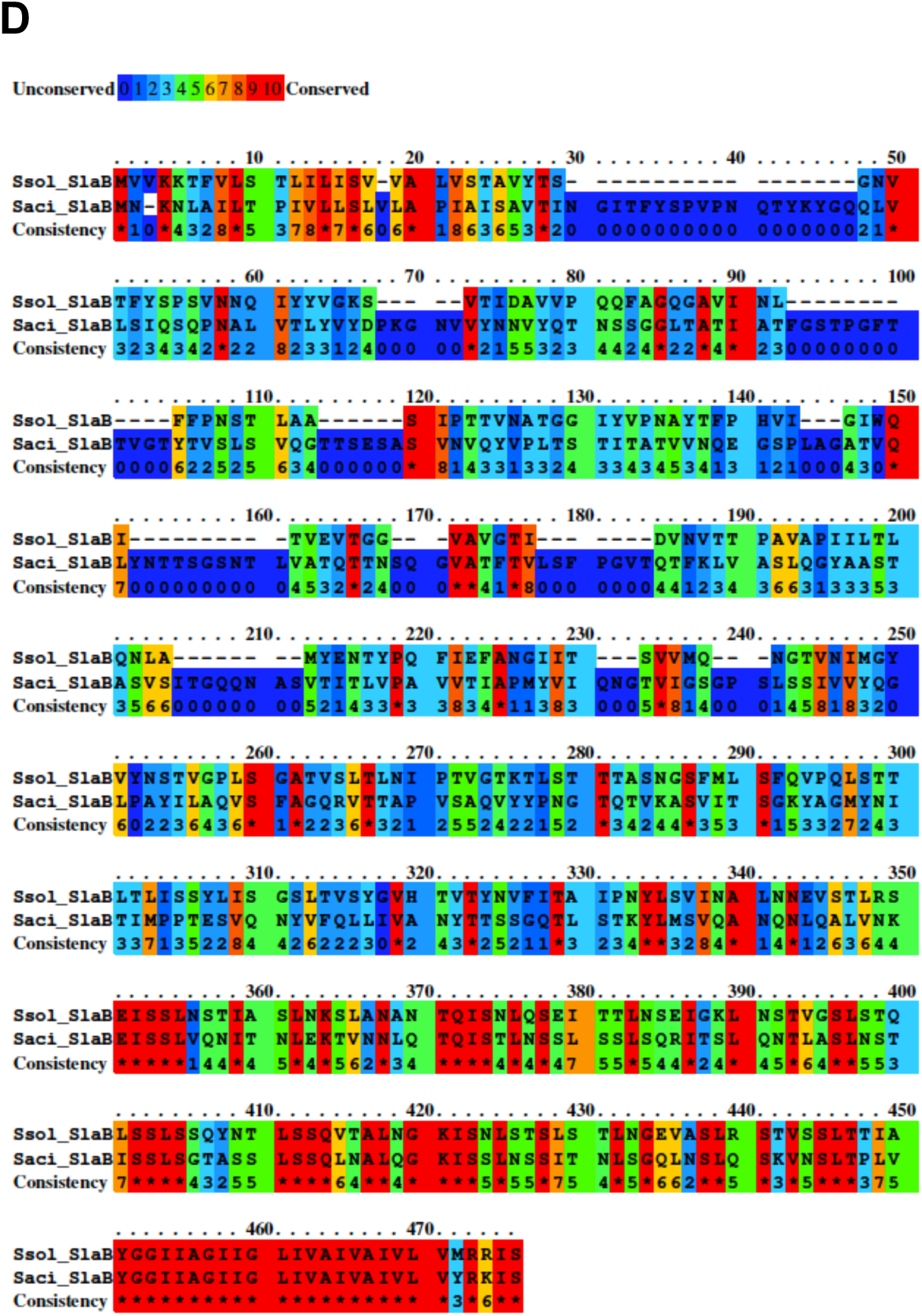
Sequence alignment comparing SlaA and SlaB from *Sisl, Ssol* and *Ssaci*. The sequence identity: *Sisl_*SlaA / S*aci_*SlaA, 24% (A); *Sisl_*SlaB / S*aci_*SlaB, 25% (B); *Ssol_*SlaA / S*aci_*SlaA, 25% (C); *Sisl_*SlaB / S*aci_*SlaB, 26% (D).

**Figure 1 S3.**
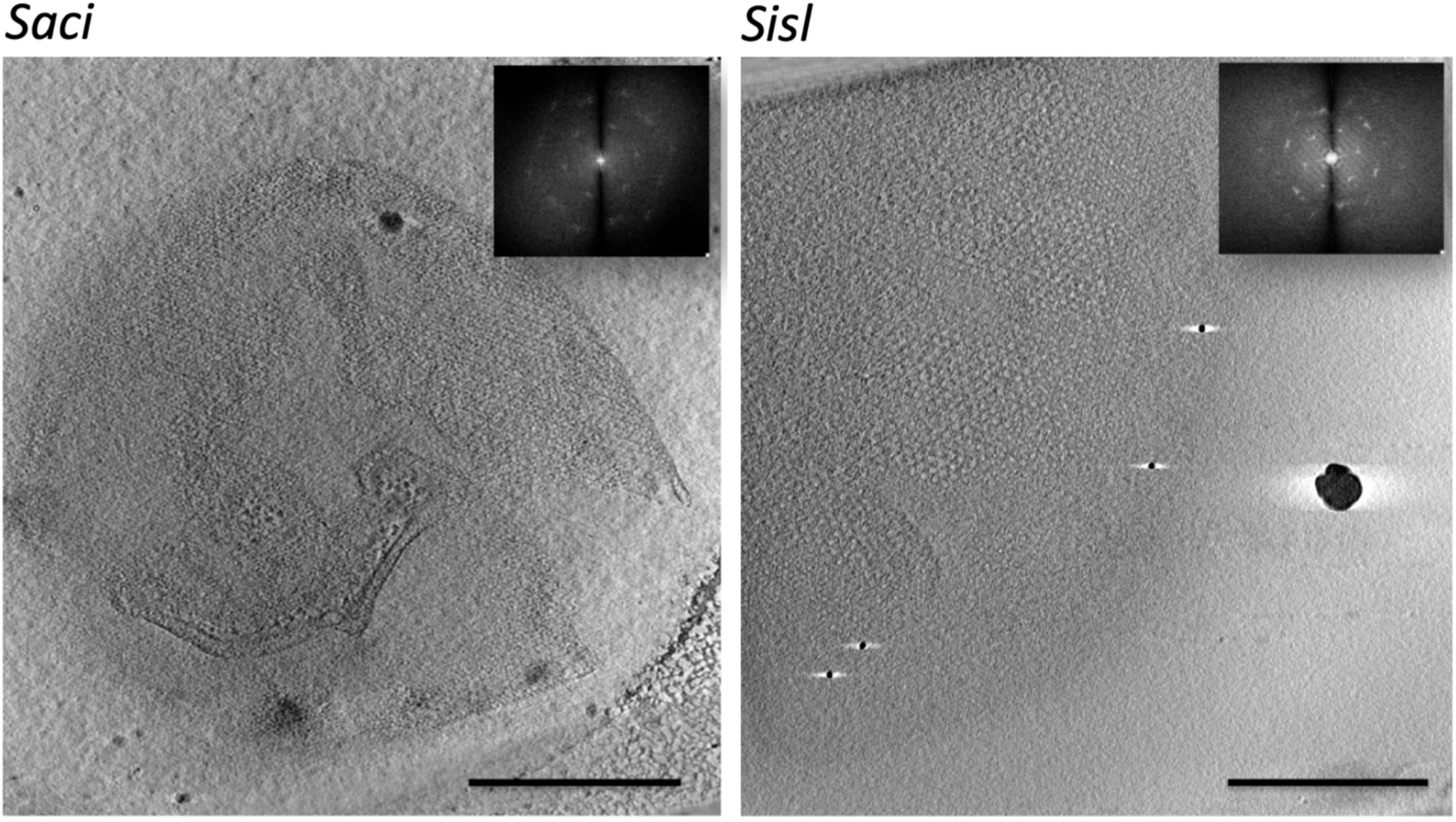
Tomographic slices through S-layer planes in whole-cell tomograms. Left panel, tomographic slice through the S-layer of Sulfolobus acidocaldarius; right panel, tomographic slice through the S-layer of Sulfolobus islandicus. Insets, power spectra reveal 2D lattices with hexagonal symmetry.

**Figure 1 S4.**
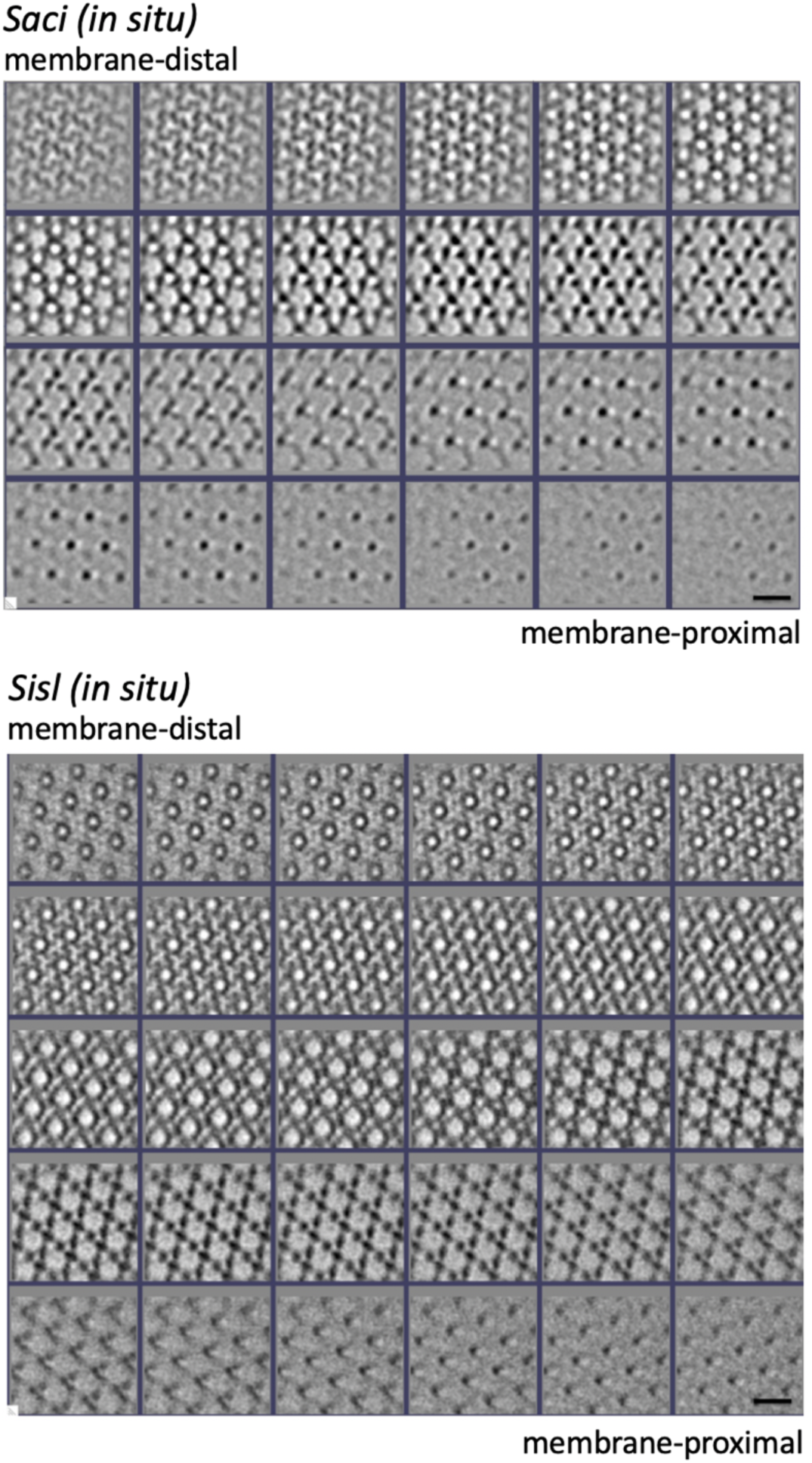
Consecutive slices through sub-tomogram averages of S-layers. Top panel, in situ map calculated from whole-cell tomogram of Sulfolobus acidocaldarius, bottom panel, map calculated from whole-cell tomogram of Sulfolobus islandicus.

**Figure 1 S5.**
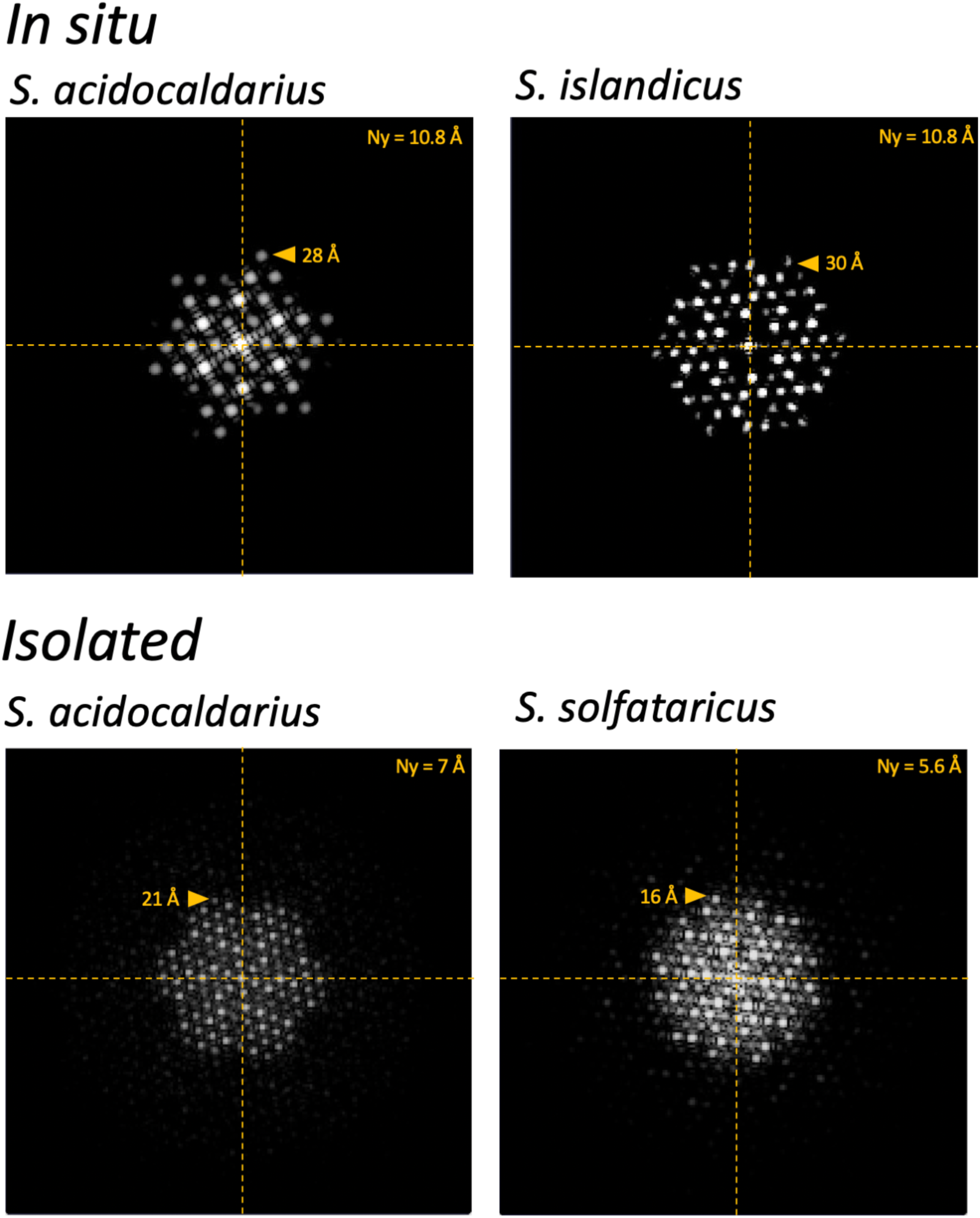
Resolution of S-layer maps determined by power spectra. Top panel, power spectra of in situ maps calculated from whole-cell tomograms, bottom panel, power spectra of maps calculated from isolated S-layers. Ny, Nyquist frequency

